# Hippocampal morphometry in sudden and unexpected death in epilepsy (SUDEP)

**DOI:** 10.1101/522300

**Authors:** Alyma Somani, Anita-Beatrix Zborovschi, Yan Liu, Smriti Patodia, Zuzanna Michalak, Sanjay M Sisodiya, Maria Thom

## Abstract

**Objective:** To determine hippocampal morphometric parameters, including granule cell dispersion and features of malrotation, as potential biomarkers for SUDEP from an archival post-mortem series.

**Methods:** In a retrospective study of 187 archival post-mortems from three groups, SUDEP (68; 14 with hippocampal sclerosis (HS)), non-SUDEP epilepsy controls (EP-C =66; 25 with HS) and non-epilepsy controls (NEC= 53), Nissl/H&E stained sections from left and right hippocampus from five coronal levels were digitised. Image analysis was carried out for granule cell layer (GCL) thickness and measurements of hippocampal dimensions (HD) for shape [width (HD1), height (HD2)] and medial hippocampal positioning in relation to the parahippocampal gyrus (PHG) length (HD3). A qualitative evaluation of hippocampal malrotational (HMAL) features, dentate gyrus invaginations (DGI) and subicular/CA1 folds (SCF) was also made.

**Results:** GCL thickness was increased in HS more than those without (p<0.001). In non-HS cases increased GCL thickness was noted in EP-C compared to NEC (p<0.05) but not between SUDEP and NEC. There was no significant difference in the frequency of DGI, SCF, measurements of hippocampal shape (HD1, HD2 or ratio) or medial positioning between SUDEP, EP-C and NEC groups, when factoring in HS, coronal level and age at death. Comparison between left and right sides within cases showed significantly greater PHG lengths (HD3) on the right side in the SUDEP group only (p=0.018)

**Conclusions:** No hippocampal morphometric features were identified in support of either excessive granule cell dispersion or features of HMAL as biomarkers for SUDEP. Asymmetries in PHG measurements in SUDEP warrant further investigation as they may indicate abnormal central autonomic networks.

## INTRODUCTION

Sudden and Unexpected death in epilepsy (SUDEP) is the leading cause of premature death in this condition^1^. Peri-ictal central autonomic disturbance in cardio-respiratory regulation is one likely mechanism, possibly mediated through abnormal functional connections between cortical, limbic and brainstem centres^2^. The limbic system, including hippocampus and amygdala, have autonomic regulatory functions and connections with the brainstem^3, 4^; experimental stimulation studies confirm hippocampal modulation of respiratory and cardiovascular activity^5^ and electrode stimulation of the hippocampus induces apnoea in some people with epilepsy^6^. Neuropathological alteration of the hippocampus, including sclerosis and granule cell dispersion, is frequent in temporal lobe epilepsy (TLE)^7^. In sudden unexplained deaths in infancy and childhood, which share some circumstantial similarities with SUDEP, developmental anomalies of the hippocampus have been recently reported^8–10^. In MRI studies in SUDEP, increased grey matter volume in the right hippocampus and parahippocampal gyrus (PHG), compared to healthy controls, was observed^11^. A systematic neuropathology analysis of hippocampal morphology in SUDEP series has not been carried out^12^. Our objective, in a large series of post-mortem (PM) cases from SUDEP and control groups, was to seek to identify a morphometric signature for SUDEP as a potential biomarker.

## METHODS

### Cases

Slides were retrieved from the UCL Epilepsy Society Brain and Tissue Bank or obtained through *Brain UK*, Queen Square and MRC brain banks from 187 mainly adult PM cases from three groups: 68 SUDEP, 66 epilepsy non-SUDEP (Epilepsy disease controls, EP-C) and 53 non-epilepsy controls (NEC). The PM tissues were collected across four decades: 31% (2010-16), 45% (2000-09), 24% (1990-99) and 0.5% (1980-89). Following review of the records, the cause of death was also further stratified into six subgroups : SUDEP into definite-SUDEP or possible/probable-SUDEP according to published criteria^13^, EP-C into non-SUDEP epilepsy-related deaths (ERD) (for example status epilepticus) or non-seizure related causes of deaths and NEC into non-epilepsy sudden deaths (NESD, including sudden cardiac death) and other causes of death (Table 1). The presence, laterality and subtype of hippocampal sclerosis^14^ was reviewed and other neuropathology confirmed at PM, including traumatic brain injury (TBI), cerebrovascular disease (CVD) or malformation of cortical development (MCD). The clinical data extracted included the age of onset of epilepsy, seizure types and syndrome if known (Table 1) and the brain weight at PM was recorded. The cases were all consented for research and the project has ethics approval (under NRES17/SC/0573).

**Table 1.** Cases included in the study, demographics and hippocampal sections. *OTHER SUDEP = Possible, Probable SUDEP. $ Epilepsy-related death (ERD) = death directly attributable to a seizures e.g. status epilepticus but not including SUDEP. ANT = anterior ventral hippocampus (rostral to Red nucleus), BODY – hippocampus body and precise level not available), COD = cause of death, LGN = lateral geniculate nucleus, NESD = non-epilepsy sudden deaths, CVD = cerebrovascular disease (any type), Gen Sz. = generalised seizures (# information on seizure type was not available in all cases or if primary or secondary generalised seizures), ID= Idiopathic epilepsy, LRE = lesion-related epilepsy (including post-traumatic epilepsy), MCD = malformation of cortical development, Red N = level of red nucleus, POST = posterior hippocampus (caudal to LGN), TBI = traumatic brain injury, Sz = seizures, TLE = temporal lobe epilepsy, DS = Dravet syndrome. N/A = not applicable. N/S = not significant (~these comparisons were made between SUDEP and EPC groups, the remainder between all groups with non-parametric tests), N/D = no detail of block side (some cases from other brain banks were provided without indication of side). Hippocampal sclerosis (HS) was subtyped according to the ILAE criteria (see text).

| GROUP<br>N = cases<br>S = sections | Subgroup<br>(N=cases<br>S=Hippocampal sections) | GENDER<br>M: F ratio | AGE DEATH<br>(range)<br>years | AGE SEIZURE ONSET<br>(range) years | Epilepsy Syndrome | Gen. Sz.<br>(% of cases<sup>#</sup>) | Partial or focal Sz.<br>(% of cases<sup>#</sup>) | Hemi-spheres<br>( Number of sections) | Coronal<br>Levels<br>(Number of Sections) | Hippocampal<br>Sclerosis<br>(Number of Sections of<br>subtypes) | Other Pathology<br>(Number of cases)<br>Brain weight mean (range, g) |
| --- | --- | --- | --- | --- | --- | --- | --- | --- | --- | --- | --- |
| SUDEP<br>N=68<br>S = 143 | DEFINITE SUDEP<br>N= 25<br>S = 44 | 62 : 38 | 38<br>(2-79) | 10.4<br>(0.5-54) | ID (9)<br>LRE (57)<br>TLE (16)<br>DS (2) | 98% | 81% | Left (65)<br>Right<br>(63)<br>N/D (15) | ANT (17)<br>RED N<br>(28)<br>BODY (20)<br>LGN (74)<br>POST (4) | N= 14<br>(Bilateral, 3<br>Right side<br>40% )<br>Type 1 (16)<br>Type 2 (2)<br>Type 3 (2)<br>No HS (123)<br>Bilateral (7) | TBI (12)<br>CVD (11)<br>MCD (8)<br>1342<br>(1005-1690 g) |
|  | SUDEP-OTHER*<br>N= 43<br>S = 99 |  |  |  |  |  |  |  |  |  |  |
| EPILEPSY CONTROLS<br>(EP-C) N=66, S=131 | EPILEPSY-RELATED DEATH<br>(ERD)<sup>§</sup><br>N= 9<br>S = 23 | 66 : 33 | 62.8<br>(20-96) | 14.4<br>(0.3-84) | ID (13)<br>LRE (76)<br>TLE (10)<br>DS (4) | 95% | 72% | Left (59)<br>Right<br>(59)<br>N/D (13) | ANT (16)<br>RED N<br>(16)<br>BODY (40)<br>LGN (55)<br>POST (4) | N=25<br>(Bilateral,<br>15<br>Right side<br>50% )<br>Type 1 (33)<br>Type 2 (3)<br>Type 3 (8)<br>No HS (87) | TBI (19)<br>CVD (31)<br>MCD (13)<br>1221<br>(788-1993<br>g) |
|  | Non-ERD<br>N= 57<br>S = 108 |  |  |  |  |  |  |  |  |  |  |
| NON-EPILEPSY<br>CONTROLS(NEC)<br>N=3, S = 75 | NESD<br>N= 13<br>S = 21 | 45 : 54 | 53<br>(23-93) | N/A | N/A | N/A | N/A | Left (22)<br>Right<br>(23)<br>N/D<br>(30)* | ANT (15)<br>RED N (6)<br>BODY (9)<br>LGN (45) | N/A | TBI (1)<br>CVD (10)<br>MCD (0)<br>1327<br>(1070-1694 g) |
|  | Non-sudden<br>death controls<br>N= 40<br>S = 54 |  |  |  |  |  |  |  |  |  |  |
| Significance<br>between<br>groups~ |  | P =<br>0.025 | P< 0.001 | N/S | P<<br>0.001~ |  |  | P< 0.001 | N/S | P< 0.001<br>(HS type) | P= 0.01-0.001 |

A total of 349 Cresyl Violet (CV) (or H&E if CV not available) hippocampal sections were retrieved from the archives (up to four sections per case). In 122 cases, paired left and right sides were included and in 52 cases more than one coronal level of hippocampus was studied. The coronal levels were grouped and recorded as : (i) anterior/pes, (ii) level of the red nucleus, (iii) hippocampal body, (iv) the level of the lateral geniculate nucleus (LGN) and (iii) posterior hippocampus. All sections were scanned on a SCN400F digital slide scanner (Leica Microsystems, Wetzlar, Germany) at 40x magnification. We excluded cases from the study with large slides (mainly prior to 2000), those with poor section quality that compromised morphometry or cases with incomplete regions of interest.

### Hippocampus shape and position measurements

On digital images, measurements of the hippocampus and parahippocampal gyrus were made at x 0.58 magnification using Leica Slidepath software (Leica Microsystems (UK), Milton Keynes, UK) and guided by published MRI studies that have evaluated hippocampal malrotation (HMAL)^15, 16^. Compared to MRI, however, analysis of tissue blocks is compromised by the lack of a perpendicular/midline reference to evaluate hippocampal axis rotation and also the collateral sulcus is not always present in small blocks. **Hippocampal size and shape (HD1 and HD2)**: Hippocampal Dimension (HD) 1 was the distance from the bottom edge of the ventricle to the subpial margin of the hippocampal sulcus (Figure 1a), both representing anatomically consistent structures between cases. HD2 was then drawn as perpendicular to HD1, spanning the maximal distance from the ventricular border adjacent to CA2 to the subiculum/white matter border (Figure 1b). The ratio of HD1 to HD2 was then calculated to give a measure of the hippocampal shape from oval (HD1/2 >1), round (HD1/2 =1) to upright (HD1/2<1), as in MRI studies of hippocampal morphology^15^. **Hippocampal medial positioning (HD3 and HD4)**: These measurements were also based on MRI studies^15^, to estimate the amount of subiculum not covered by hippocampus as one criterion in the evaluation of HMAL (Figure 1a). These measurements were taken parallel to each other: HD3 was maximum distance measured along the main axis of the subiculum extending from the medial ventricular border of the hippocampus to the outer edge of the parahippocampal gyrus (PHG). HD4 was then measured parallel to HD3 from the ventricle to the opposite edge of the hippocampus body. The ratio of HD4/3 then gave an estimate of the relative medial position of the hippocampus. In anterior/pes hippocampus sections with double representation of the dentate gyrus, the measurement is taken from the medial hippocampus part only (Figure 1b). Repeated measurements for were carried out on 82 cases with good reliability (intraclass correlation coefficient = 0.9, p<0.0001).

**Figure 1.**
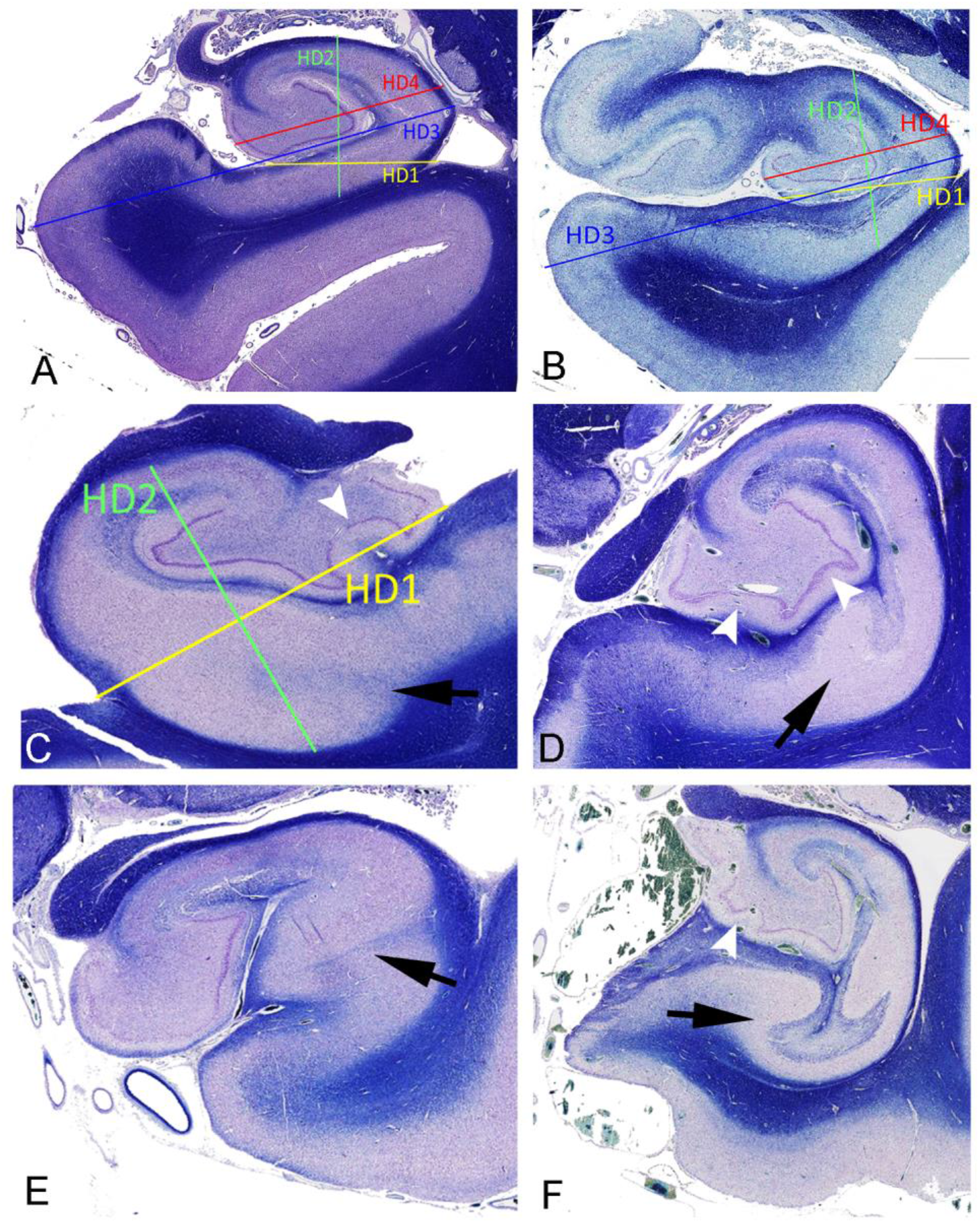
Illustration of hippocampal shape and position measurements, subiculum/CA1 folds and dentate gyrus invagination. A. Section of hippocampus at level of lateral geniculate nucleus (LGN). Hippocampal Dimension (HD) 1 (yellow line) was a line from the bottom border of the ventricle to the subpial surface of the hippocampal sulcus. HD2 (green line) was then drawn as the widest perpendicular to HD1, spanning from the ventricular border adjacent to CA2 to the subiculum/white matter border (Figure 1b). The ratio of HD1 to HD2 provide a measure of **hippocampal shape** from oval to round. HD3 (blue line) and HD4 (red line) were taken parallel to each other; HD3 is the maximal length from the medial ventricle border of the hippocampus to the outer edge of the parahippocampal gyrus (PHG) running alongside the length of the subiculum. HD4 measurement was then taken from the ventricle border to opposite edge of the hippocampus body and. HD3 and HD4 ratio gives an approximation of **hippocampal medial positioning** as based on MRI studies^15^ to estimate the amount of subiculum not covered by hippocampus as one criteria in the evaluation of hippocampal malrotation (HMAL). B. An anterior hippocampus sections with double representation of the dentate gyrus where the measurements were made from the medial part only. **C to F** show examples from different cases with different severity of HMAL with folds or bulbous expansions in the subiculum/CA1 segment (arrow) and invaginations in the dentate gyrus (white arrowheads). **C** is from a non-epilepsy control (HD1 and HD2 measurements are also superimposed as an illustration), **D** and **F** from an epilepsy-control with bilateral hippocampal changes. All shown in Cresyl violet/LFB stain from the original magnification of x 0.58 on the Leica slidepath software (Leica Microsystems (UK), Milton Keynes, UK) at which the measurements were taken (C-D further magnified to show detail).

### Granule cell layer (GCL)

We carried out qualitative evaluation of the GCL based on the dominant pattern in both dentate blades: normal and compact GCL (Pattern 1), compact GCL but appears broader (Pattern 2), unequivocal GCL dispersion (Pattern 3), mild or focal GCL dispersion (Pattern 4) and granule cell depletion (Pattern 5) (Figure 2a-e). We then measured the GCL thickness on the external blade (adjacent to CA2, GCL_E_) and internal blade (adjacent to subiculum, GCL_I_) of the dentate gyrus in each section. For each blade, 4 to 5 images were grabbed at x 20 magnification. Using image pro plus, the ‘best-fit’ basal line was drawn using the 10 deepest granule cells at the hilar border. This basal line was used to measure the perpendicular distance of the 10 outermost granule cells in each image (Figure 2f). The average of these provided a mean GCL_E_ and GCL_I_ thickness as based on the extent of maximal cell dispersion; the mean of these two values (GCL_M_) was also calculated for each slide. GCL measurements were repeated in 10 randomly-selected slides with good intra-observer reproducibility (intraclass correlation coefficient = 0.9, p<0.0001; mean of differences 2.7microns, SD = 11.07 microns).

**Figure 2.**
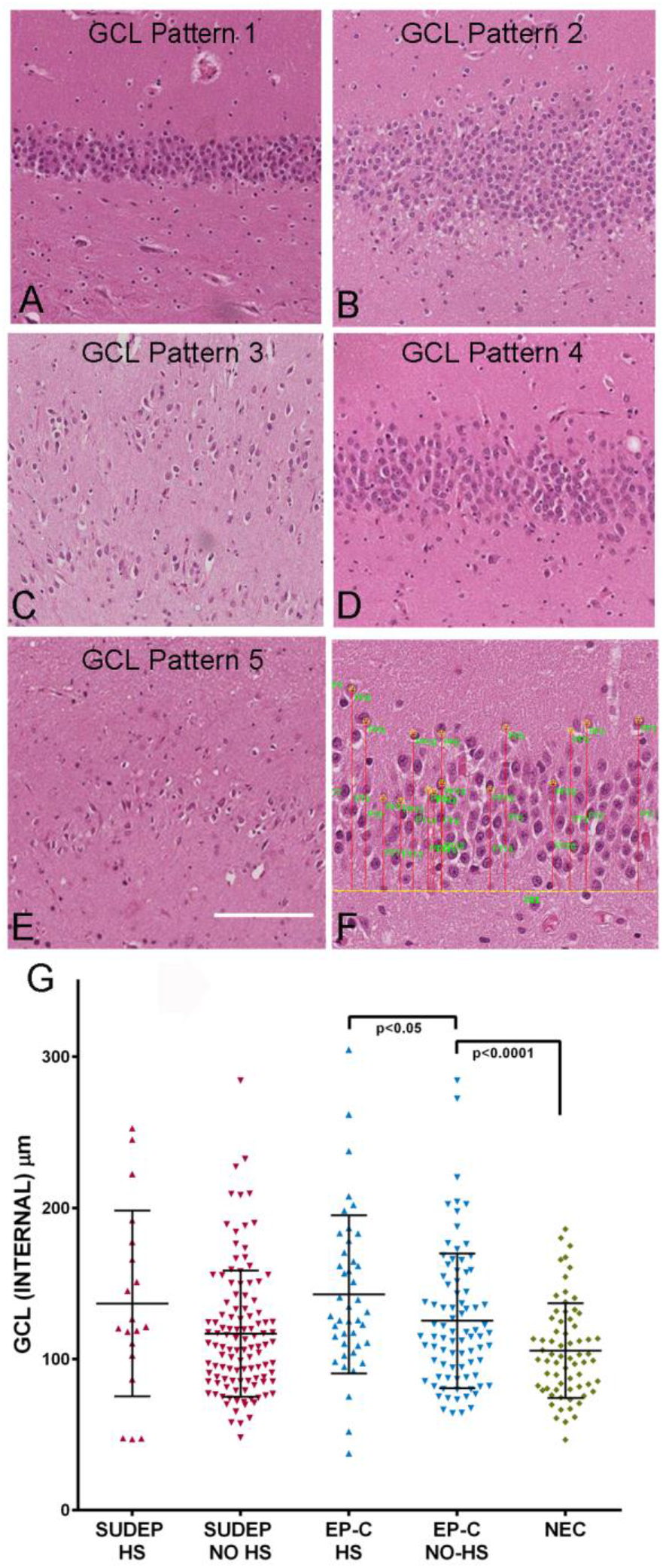
Granule cell layer patterns and measurements. **A-E** Illustrations of the five qualitative patterns in the granule cell layer (GCL) as detail in text. In brief : Pattern 1 (A) normal compact GCL, Pattern 2 (B) mild broadening of the GCL, Pattern 3 (C) clear dispersion of cells in GCL, Pattern 4 (D) mild, focal dispersion in GCL and Pattern 5 (E) loss of granule cells as the predominant finding. F. Illustrates the method used for measurement of GCL width with ‘best fit’ basal cell layer line shown in yellow from which the perpendicular distances of the 10 most distal granule cells were measured (red lines). G. Scatter graph of measurements for GCL thickness measured in microns in the internal blade of the dentate gyrus between groups of SUDEP, epilepsy controls (EP-C) and non-epilepsy controls (NEC) are shown. The epilepsy groups have also been split for the presence of hippocampal sclerosis (HS) showing that GCD was greater in HS than non-HS cases, with significant differences in EP-C group (but not SUDEP). There was significantly higher GCD measurements in non-HS EP-C cases than NEC but this was not observed in the SUDEP group. White bar in A-E equivalent to 100 microns.

### Qualitative hippocampal patterns

The presence of invaginations along the straight sections of the DG, on either the CA2 side or SC side, was recorded as present or not (Figure 1c-e). The presence of folds in the CA1/subiculum pyramidal cell layer was also recorded (Figure 1c-d).

### Statistical methods

Measurements were compared between groups using non-parametric tests and corrected for multiple comparisons. The Wilcoxon test was used for comparison of paired hemispheric samples. Spearman’s correlation test was used for correlation of clinical factors, such as age, with hippocampal measurements. Multiple regression analysis was used to evaluate the influence of multiple factors (including the presence of sclerosis, anatomic level, age and cause of death) on hippocampal measurements. Values of p≤0.01 were taken as significant.

#### Data availability policy

Data will be made available at the request of other investigators for purposes of replicating procedures and results.

## RESULTS

### Cases

60% of the cases were male with a mean age at death of 50 years (2-96 years); there were some group differences, with more males in the epilepsy groups and a significantly younger age of death in the SUDEP group (Table 1). HS was confirmed on re-examination of the sections in 14 SUDEP and 25 EP-C; HS was bilateral in 18 cases, and showed type 1 HS pattern in 32 cases. HS involved the right side in 40% of SUDEP and 50% in EP-C groups (Table 1).

### Hippocampal dimensions, shape and size

Hippocampal dimensions HD1, HD2 and were significantly lower in HS (all types) than No-HS cases (p<0.0005) (Figure 3a, Table 2). HD1/HD2 ratio was also lower in HS cases than No-HS (Figure e-1a) indicating a more rounded than elliptical shape. There were also significant differences in these measures (HD1, HD2 and its ratio) between HS subtypes (p<0.01 to <0.0001) (Figure 3c). HD1 and HD2 positively correlated with brain weight in all cases (p<0.01) except cases with HS. There was significant variation in HD1, HD2 and HD1/HD2 ratio with the coronal level of the hippocampus in both HS and non-HS cases (p<0.01 to 0.0001) (Figure 3b).

**Figure 3.**
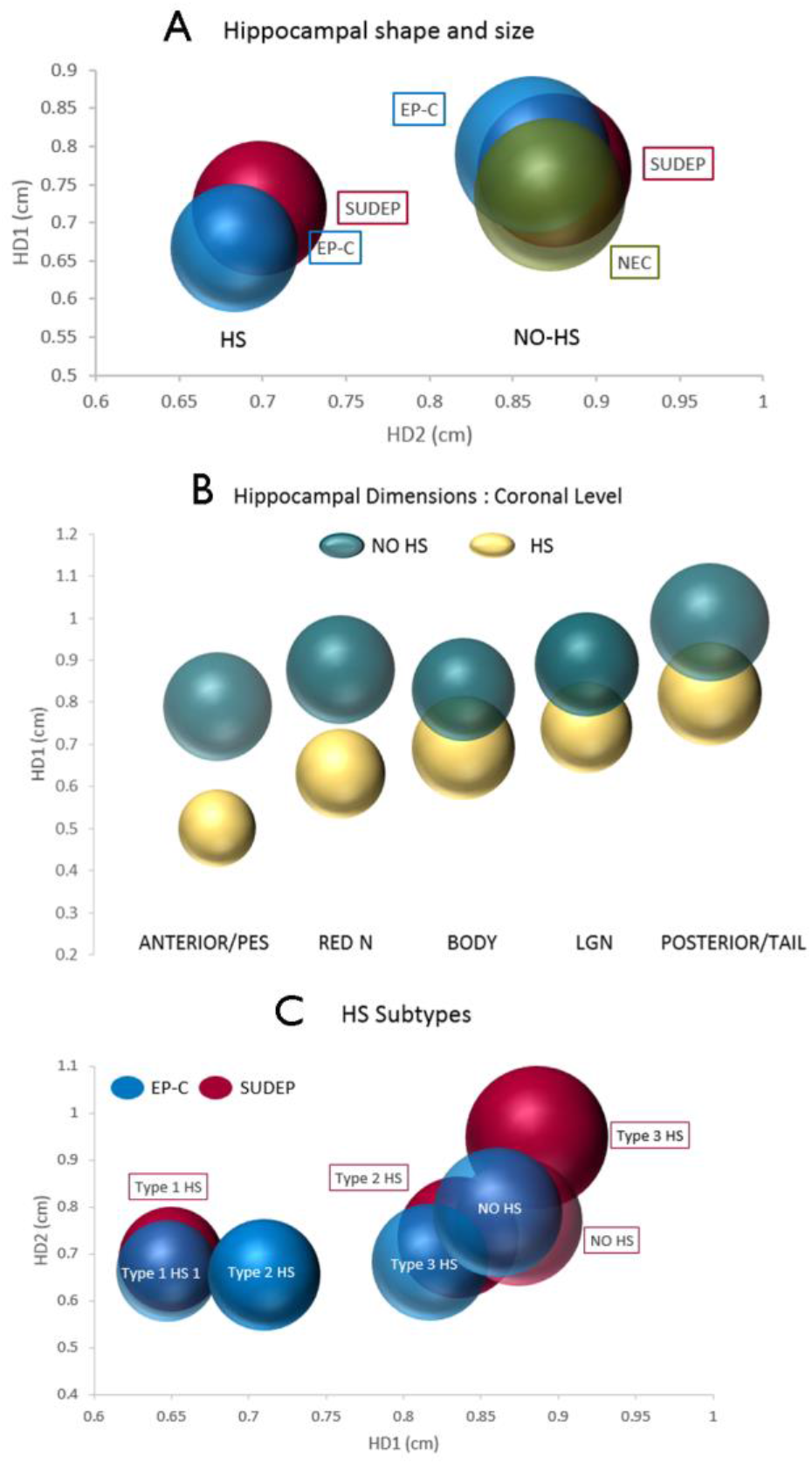
Hippocampal dimensions and shape in SUDEP and control groups. A. Bubble graph of hippocampal dimension (HD)1 plotted against HD2 in the three cause of death groups (SUDEP (red), Epilepsy-controls (EP-C, Blue)) and non-epilepsy controls (NEC, Green) groups splitting hippocampal sclerosis (HS) cases from No-HS cases. These represent the mean values in each group and the size of bubble is equivalent to relative mean area of the hippocampus based on HD1 and HD2 measurements (for graphical representation and comparison, shown as spheres rather than ellipses). There were significant differences between HD1 and HD2 between HS and No-HS cases; no significant differences in HD2 and HD1 or ratio (HD1/HD2) between cause of death groups was shown in No-HS groups. (In on line figure e1a scatter graphs of the ratio HD1/HD2 in these groups is shown. B. Bubble graphical representation of the variation of HD1 with coronal level of the hippocampus taken from one of five levels as detailed in the text : (i) anterior/pes, (ii) level of the red nucleus (RED N), (iii) hippocampal body, (iv) the lateral geniculate nucleus level (LGN) and (iii) posterior hippocampus or tail. The cases are split into HS and No HS cases and the area of the bubble represents the relative mean area of the hippocampus based on HD1 and HD2 measurements. There was a significant variation in the HD1 (as well as HD2 measures and its ratio, data not shown) between five coronal levels for both HS and No-HS cases (p<0.01). C. Bubble graphical representation of HD2 between Hippocampal subtypes (ILAE types 1, 2 and 3) shown in both SUDEP and EP-C groups. The area of the bubble represents the relative mean area of the hippocampus based on HD1 and HD2 measurements. Significant variation in HD2 (as well as HD1 and HD1/HD2 ratio, data not shown) was observed between HS subtypes (p<0.01) but these variations were not significantly different between SUDEP and EP-C groups.

**Table 2.** Results of Hippocampal Measurements (hippocampal dimensions HD1, 2, 3 and 4 : see text for details) and Granule cell layer measurements (GCL) in the three main groups, SUDEP, Epilepsy controls and non-epilepsy controls with mean values and Standard deviations (SD) shown.

|  | **HD1 (cm)**<br>HS (SD)<br>NON-HS<br>(SD)<br>N=case<br>number | **HD2 (cm)**<br>HS (SD)<br>NON-HS<br>(SD)<br>N=case<br>number | **HD3 (cm)**<br>HS (SD)<br>NON-HS<br>(SD)<br>N=case<br>number | **HD4 (cm)**<br>HS (SD)<br>NON-HS<br>(SD)<br>N=case<br>number | **GCL**<br>**EXTERNAL**<br>**(μm)**<br>HS (SD)<br>NON-HS (SD)<br>N=case<br>number | **GCL**<br>**INTERNAL**<br>**(μm)**<br>HS (SD)<br>NON-HS (SD)<br>N=case<br>number |
| --- | --- | --- | --- | --- | --- | --- |
| SUDEP | 0.69 (.15)<br>0.87 (.18)<br>N= 138 | 0.71 (.14)<br>0.76 (.13)<br>N= 138 | 1.6 (.22)<br>1.82 (.27)<br>N=132 | 0.73 (.14)<br>0.91 (.14)<br>N=132 | 143 (79)<br>109 (35)<br>N= 135 | 136 (61)<br>116 (41)<br>N=135 |
| EPILEPSY<br>CONTROLS | 0.68 (.25)<br>0.86 (.21)<br>N= 123 | 0.66 (.13)<br>0.78 (.17)<br>N= 123 | 1.46 (.22)<br>1.72 (.29)<br>N=119 | 0.69 (.21)<br>0.91 (.18)<br>N=119 | 131 (56)<br>119 (33)<br>N=125 | 142 (52)<br>125 (44)<br>N=125 |
| NON<br>EPILEPSY<br>CONTROLS | -<br>0.87 (.19)<br>N=70 | -<br>0.73 (.15)<br>N=70 | -<br>1.72 (.29)<br>N=68 | -<br>0.91 (.19)<br>N=68 | -<br>104 (24)<br>N=69 | -<br>105 (3)<br>N=69 |

#### HD1/HD2 between case groups

There were no significant differences in HD1, HD2 measurements and HD1/HD2 ratio between SUDEP, EP-C and NEC groups when the presence of HS and the anatomical coronal level was accounted for with multivariate analysis (Figure 3a, Figure e-1a, Table 2). When the causes of death were further stratified into six groups, including Definite-SUDEP, there were also no significant differences in HD1, HD2 and HD1/HD2 between the groups. There were also no significant difference in measurements between HS subtypes in SUDEP and EP-C groups (Figure 3c).

### Hippocampal medial positioning

Hippocampal dimension HD3 (PHG length), HD4 (hippocampal body) and its ratio HD4/HD3 (relative medial position of the hippocampus) were all significantly lower in HS (all types) than No-HS cases (p<0.0001) (Table 2). HD3 also significantly correlated with brain weight (p<0.0001) in No-HS but not HS cases. There was significant variation in HD3, HD4 and HD4/HD3 ratio relative to the coronal level of the hippocampus in all cases (p<0.01 to 0.005) as well as for no-HS cases for HD3 and HD4/HD3 (p<0.01 to 0.005) (Figure e-1b).

#### HD3/HD4 between case groups

There was significant variation noted in HD3 between SUDEP, EP-C and NEC groups (p<0.0005), including analysis of No-HS cases alone (p=0.01), with greater mean lengths for the parahippocampal gyrus (HD3) in the SUDEP group (Table 1). Significant variation in HD3 was also noted when causes of death were sub-classified into the six subgroups, for all cases and No-HS cases (p<0.0005) with highest values in the Definite-SUDEP group (Figure 4a). The significance of HD3 measurement differences between the cause of death groups diminished when factoring for different coronal levels (p<0.03). Also, the SUDEP group had significantly younger mean age at death than control groups; when this was factored using multivariate analysis, no significant differences were noted for HD3 between all the cause of death groups.

**Figure 4.**
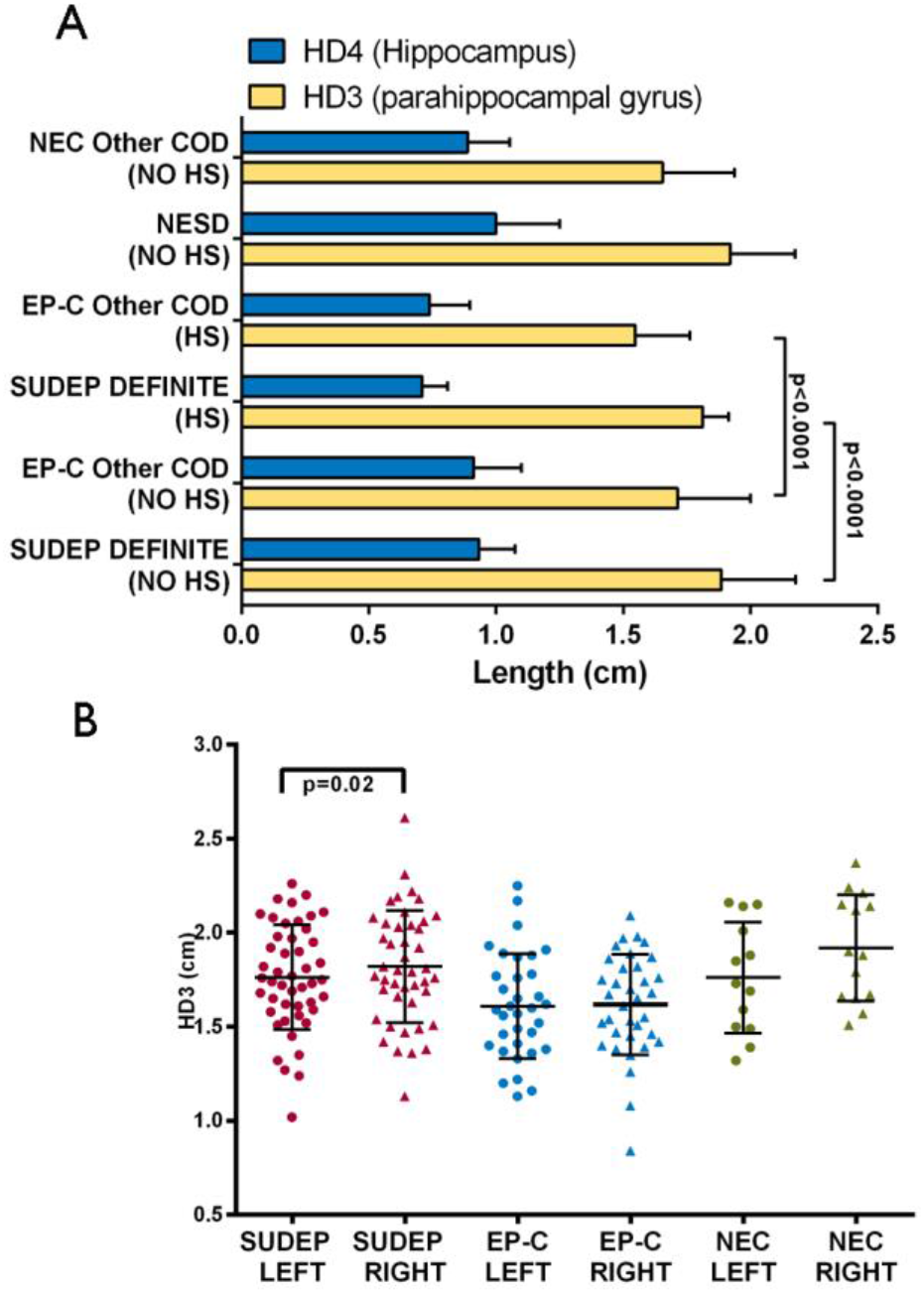
Hippocampal measurements for medial position of the hippocampus. A. Bar chart showing mean values (error bars are standard deviation) for hippocampal dimensions HD3 (Parahippocampal gyrus) and HD4 (hippocampus) (see text for details) in cause of death groups : (i) SUDEP-definite, (ii) Epilepsy-controls (EP-C) with cause of death other than epilepsy-related, (iii) non-epilepsy sudden deaths (NESD) and (iv) other non-epilepsy controls (NEC). The epilepsy groups are further divided into cases with or without HS. HD3 and HD4 were significantly lower in HS cases than non-HS. There were no significant differences for the medial positioning of the hippocampus (HD4 relative to HD3) between cause of death groups. HD3 measurement was highest in SUDEP cases, but when the age of the patient and coronal level was factored, this difference was not significant. B. Scatter graph of HD3 measurements in cases with paired left-right hippocampal sections in SUDEP, EP-C and NEC groups. The right HD3 was greater than left in all groups but reaching significance only in the SUDEP group.

### GCL measurements

Qualitative evaluation of GCL patterns was undertaken in 244 sections from the SUDEP and control groups (Table 3); significant differences in GCL_M_ measurements between the GCL patterns were noted, as anticipated, with highest values in cases with dispersion, lowest values in cases with granule cell depletion and intermediate values with focal or mild dispersion (Kruskall Wallis p<0.0001) (Table 3, Figure e-1c). This supported the assertion that our quantitative measures reflect qualitative patterns reported in previous similar studies^8^. Greater values for GCL_E_, GCL_I_ and GCL_M_ were present in HS than No-HS cases (p < 0.005 to 0.0001) (Table 2). There was also a significant difference noted for GCL_M_ between HS subtype (p≤0.005) (Figure e-1d) but not for the coronal level of the hippocampus.

**Table 3.**
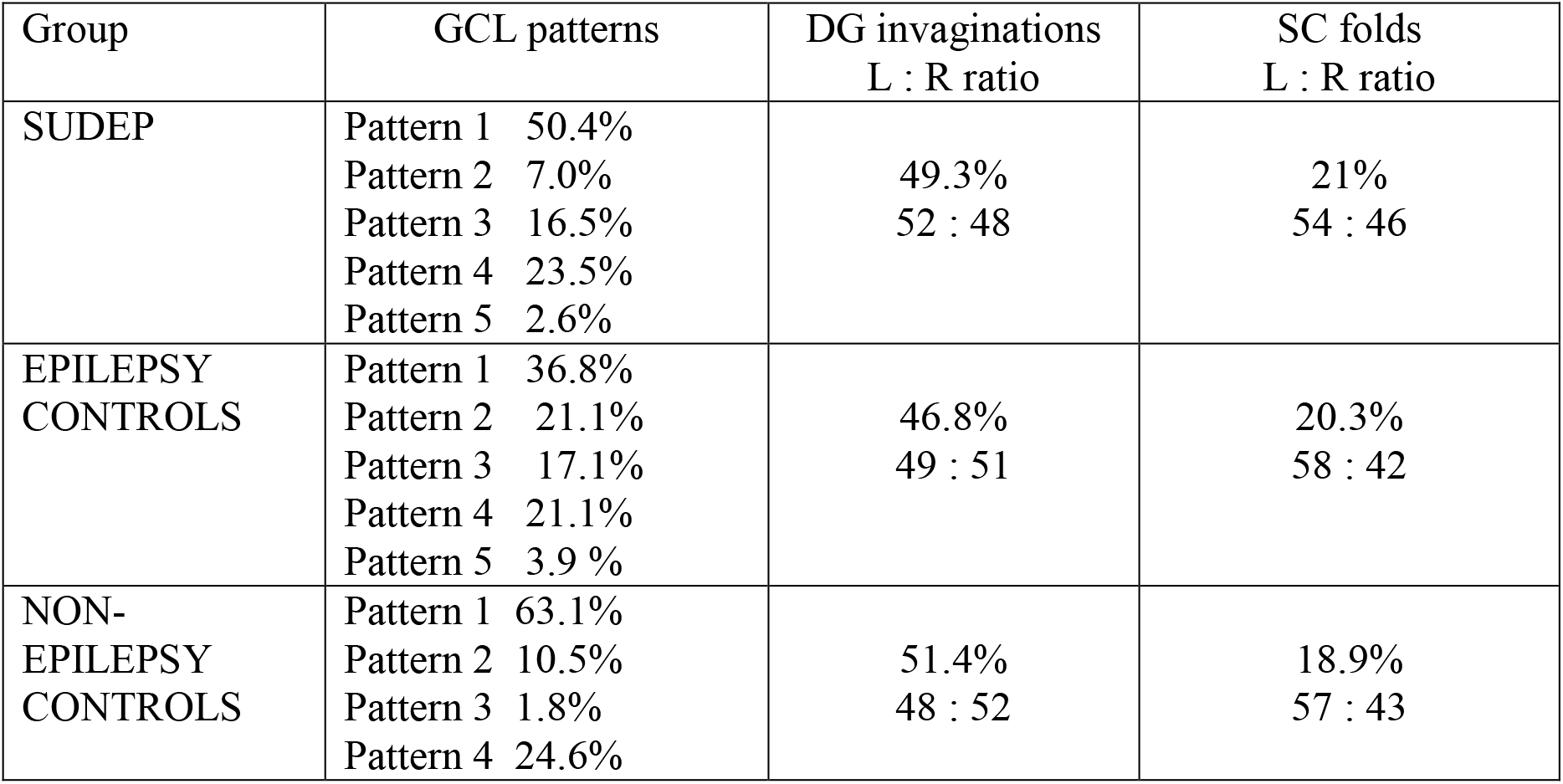
Qualitative evaluations of the frequency of hippocampal morphological features in the three cause of death groups including granule cell layer (GCL) patterns, the presence of dentate gyrus (DG) invaginations and subiculum (SC) folds. There were no significant differences in the presence of these features and the cause of death groups.

| Group | GCL patterns | DG invaginations<br>L : R ratio | SC folds<br>L : R ratio |
| --- | --- | --- | --- |
| SUDEP | Pattern 1 50.4%<br>Pattern 2 7.0%<br>Pattern 3 16.5%<br>Pattern 4 23.5%<br>Pattern 5 2.6% | 49.3%<br>52 : 48 | 21%<br>54 : 46 |
| EPILEPSY<br>CONTROLS | Pattern 1 36.8%<br>Pattern 2 21.1%<br>Pattern 3 17.1%<br>Pattern 4 21.1%<br>Pattern 5 3.9 % | 46.8%<br>49 : 51 | 20.3%<br>58 : 42 |
| NON-<br>EPILEPSY<br>CONTROLS | Pattern 1 63.1%<br>Pattern 2 10.5%<br>Pattern 3 1.8%<br>Pattern 4 24.6% | 51.4%<br>48 : 52 | 18.9%<br>57 : 43 |

#### GCL between case groups

Between groups in No-HS cases, GCL measurements were significantly increased in EP-C compared to NEC (GCL_M_, p<0.0001) but not between SUDEP and NEC (Figure 2g, Table 2). When cause of death was sub-classified into six subgroups, higher GCL_I_ was still noted in the non-ERD EP-C than both non-epilepsy control groups (p<0.05) whereas neither SUDEP group was significantly different from controls.

### SC folds and DG invaginations

There was a positive correlation between the presence of excessive folds in the subiculum and invaginations in the dentate gyrus across all cases (p = 0.004). There was no relationship between either of these features and the cause of death groups, the presence of HS, or the coronal level of the hippocampus or side (Table 3).

### Laterality

From 122 cases with paired samples from left and right hippocampus, there was alignment for the same coronal level in 88 cases. Measurements of HD1, HD3 and HD4 were higher on the right side with significant differences noted for HD3 in all pairs (p=0.016), only coronal-aligned paired sections (p=0.012) and No-HS cases alone (p=0.007). Within groups, HD3 was noted to be greater on the right side in SUDEP (p=0.018) but not in EP-C or NEC control groups (Figure 4). There were no significant differences observed for GCL measures between any groups in relation to laterality.

### Clinical-pathology correlations

Epilepsy syndromic diagnosis or seizure type was not available in a proportion of SUDEP and EP-C cases (Table 1). Where seizure type was recorded, the majority of cases in SUDEP and EP-C groups had either primary or secondary generalised and focal seizures (Table 1) but there was no significant difference in age of seizure onset between groups. There was a positive correlation between the age of onset of seizures and GCL_M_ (p<0.05) in EP-C groups but not for HD measures in any group. There was no association between GCL or HD measurements and the presence of additional TBI, CVD or MCD pathology at PM. There was a significant negative correlation between age at death and HD1, HD2, HD3 and HD4 in all cases (P < 0.001) as well as the subset of HS cases (p<0.05 to 0.005). There was no correlation with GCL thickness and age of death in any group.

## DISCUSSION

In a retrospective study of 68 SUDEP cases, we did not find evidence for differences in hippocampal position or shape or abnormalities of the granule cell layer compared to control groups after consideration of the presence of HS, the anatomical level of the hippocampus and the patient’s age, which all independently influence these features. We identified increased PHG gyrus asymmetry in SUDEP, in keeping with previous MRI studies showing increased right PHG volumes in SUDEP^11^, which may be of relevance to central autonomic pathway dysfunction.

This was a retrospective study and we aimed to maximise the number of cases to increase its power, using similar protocols to previous large published PM studies of hippocampal morphology^8^, but implementing additional morphometric quantitative analysis to reduce observer bias. The study included cases from different tissue banks acquired over different eras with inherent variations in tissue fixation and laboratory processing protocols, all of which could differentially influence tissue volume and effect measurements. It is less likely, however, that these factors affect the comparisons of measurements between hemispheres within cases or ratio of measurements used (HD1/HD2 and HD4/HD3). Further limitations were that we did not have sections from both hemispheres at multiple coronal levels in all cases or macroscopic or MRI images to enable comparison with histological measurements.

### Diminished granule cell migration in SUDEP

Granule cell dispersion is commonly associated with HS in TLE, noted in 76% of surgical specimens in a recent series^17^. In the context of HS, seizure-induced loss of reelin-synthesising interneurons has been shown to initiate neo-migration of mature granule cells^18^. Granule cell abnormalities have also recently been recognised in the context of unexplained deaths in childhood and infancy without evidence of HS^10^. In sudden unexplained death in childhood (SUDC), dentate gyrus abnormalities and HMAL were associated with a history of febrile seizures^9 19^. In sudden infant death syndrome (SIDS), focal bilamination of the dentate gyrus was identified in 41% of 114 cases^8^. It was suggested these developmental anomalies represented disease biomarkers and could be mechanistically linked to cause of death through abnormal limbic-brainstem connections that regulate cardio-respiratory control. SUDEP has parallels with SUDC/SIDS, including circumstances surrounding death, with frequent prone body position and nocturnal occurrence, which may indicate pathophysiological similarities. Indeed, it has been proposed that some SIDS/SUDC represent seizure-related deaths^20, 21^.

Dentate granule cell dispersion has been anecdotally recorded in 4% of SUDEP PM reports^12, 22^, but not systematically evaluated through review of histology sections. In this current retrospective study, based on a similar archival review protocol to the published SIDS studies^8^, but incorporating quantitative rather than qualitative evaluation, GCL measurements did not reveal excessive dispersion or broadening in SUDEP. We noted anticipated differences in GCL thickness in relation to HS type and age of seizure onset, in keeping with observations in TLE^23^. In non-HS cases however, significant GCL broadening was only observed between the non-SUDEP epilepsy group and non-epilepsy controls. This may paradoxically implicate a deficit in seizure-mediated granule cellular migratory mechanisms operating in SUDEP.

### Hippocampal malrotation is not over-represented in SUDEP

HMAL (or incomplete hippocampal inversion) has been implicated as an underlying pathological substrate for febrile seizures^24^ as well as some SUDC^10, 19^. We have previously noted severe HMAL in 9.7% of SUDEP cases, but a systematic series review is lacking^25^. Evaluation of HMAL is confounded by lack of universally applied diagnostic criteria, both for MRI and neuropathology, to enable reliable comparisons of its incidence across disease groups in published studies^15, 26^. Reported incidences of HMAL vary from 8%^27^ to 16%^16^ in non-HS TLE, 8.8% in febrile status^28^ to 88% in SUDC associated with FS^9^. Furthermore, quantitative MRI evaluations of healthy controls report an incidence of HMAL in 23-24%, more commonly involving the left side^15, 16^.

As HMAL represents a spectrum of severity, tissue pathology may be more sensitive in detecting subtle anomalies. MRI criteria include measures of hippocampal roundness and verticality, mesial positioning, subiculum thickness and depth or verticality of the collateral sulcus^15, 16, 24^. The main obstacle in the quantitative evaluation of HMAL in pathology sections is the lack of a vertical midline or adjacent sulcus as a reference^15, 16^, macroscopic images of the brain being not available in all cases, particularly for coroners’ sudden death investigations^12^. In PM cases, excessive folding of the CA1/subiculum is regarded as the most striking feature in HMAL^29^; similar ‘tectonic’ hippocampal abnormalities are described in surgical hippocampal resections, with ‘bulbous expansions’ or convolutions in the subiculum/CA1 region, typically associated with corresponding invaginations of the dentate gyrus^27^.

In this SUDEP series, our quantitative evaluation of hippocampal shape and position was based on consistently present anatomical landmarks in addition to qualitative assessment for HMAL. We noted a significant association between the presence of folds in subiculum/CA1 and dentate gyrus invaginations in keeping with previous studies^27^. Subicular/CA1 folds were present in 21% of SUDEP cases and more common on the left side but not significantly different to control groups. This, together with a lack of quantitative evidence of increased roundness or medial positioning of the hippocampus, supports the notion that, although even severe cases of HMAL may be seen in SUDEP, it is not over-represented.

### Parahippocampal gyrus asymmetry in SUDEP

We noted asymmetries in the length of the PHG along the subiculum, which was significantly greater on the right than on the left side in the SUDEP group only. Although we did not measure the volume of the PHG, this finding is in keeping with MRI observations showing increased grey matter volume in the right PHG in individuals who went on to have SUDEP compared to healthy subjects^11^. Asymmetries in the mesial temporal structure are recognised; the right hippocampus develops earlier than the left side^30^ and in healthy adults the right hippocampal volume is greater^31^. Recent data from the ENIGMA study which analysed data from over 17,000 MRIs has shown asymmetry in the cortical thickness measures in the parahippocampal gyrus (leftward asymmetry) and entorhinal cortex (rightward asymmetry) with both regions showed significant variability between sexes^32^; of note in both our epilepsy groups there was an overrepresentation of males compared to controls. Cortical asymmetries likely reflect left/right functional differences and there is increasing evidence of the role of the limbic region in autonomic regulation. For example, functional imaging studies show connections of the right hippocampus to the dorsal vagal nuclei in the regulation of heart rate after exercise^33^. Quantitative MRI studies have shown negative correlation between the volume of right PHG and heart rate variability, supporting a regulatory role of this region in parasympathetic activity^34^. The observation of PHG asymmetry in SUDEP is therefore worthy of further investigation in view of the potential influence of this region on central autonomic networks and potential dysfunction.

## Acknowledgments

We are very grateful for provision of additional SUDEP and control material for this study from the following resources: Tammaryn Lashley and the Queen Square Brain Bank at UCL London and the MRC Sudden Death Brain Bank in Edinburgh. Tissue samples were also obtained from David Hilton at Derriford Hospital as part of the UK Brain Archive Information Network (BRAIN UK) which is funded by the Medical Research Council and Brain Tumour Research.

---

**Supplemental Figure e1.**
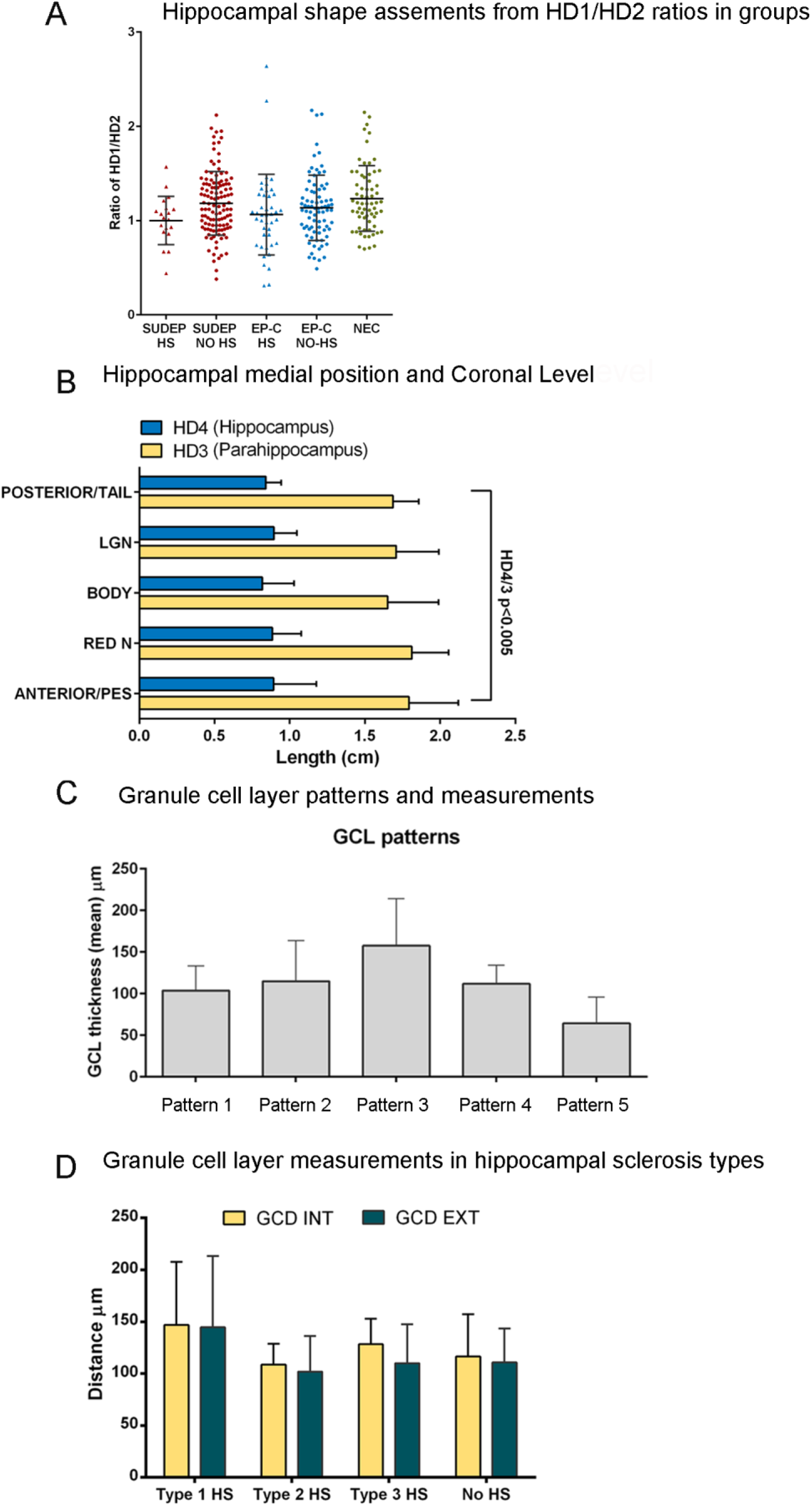
A. Scatter graph of the HD1 to HD2 ratios in each case shown for three groups (SUDEP, Epilepsy Controls (EP-C) and Non-epilepsy controls (NEC). The epilepsy groups are further divided into hippocampal sclerosis (HS) and No-HS cases. There were significantly lower HD1/HD2 ratios in Hippocampal sclerosis (HS) cases (mean 1.04, close to spherical) compared to Non-HS cases (mean 1.18, elliptical shape). B. Bar chart showing mean values (error bars are standard deviation) for hippocampal dimensions HD3 (Parahippocampal gyrus) and HD4 (hippocampus) (see text for details) in all cause of death groups with coronal level of the hippocampus taken from one of five levels as detailed in the text : (i) anterior/pes, (ii) level of the red nucleus (RED N), (iii) hippocampal body, (iv) the lateral geniculate nucleus level (LGN) and (iii) posterior hippocampus or tail. Cases are shown only for No-HS cases. There was significant variation between HD3 and HD4 measurements and coronal level (p<0.01). C. Bar chart of the mean (and standard deviation, shown as error bar) of the granule cell layer (GCL) measurements in the five qualitative patterns observed in the dentate gyrus: normal and compact GCL (Pattern 1), compact GCL but appears broader (Pattern 2), unequivocal GCL dispersion (Pattern 3), mild or focal GCL dispersion (Pattern 4) and granule cell depletion (Pattern 5). There was a significant variation in GCL measurements between the groups (p<0.0001). D. Bar chart of GCL measures in the internal (INT) and external (EXT) blade in ILAE hippocampal sclerosis subtypes. There was a significant variation noted (p<0.005) with higher values in type 1 HS cases.

## Appendix 1

| Name | Location | Role | Contribution |
| --- | --- | --- | --- |
| Alyma Somani | UCL Queen square<br>Institute of<br>Neurology | Author | Manuscript<br>preparation and<br>data acquisition |
| Anita-Beatrix<br>Zborovschi | UCL Queen square<br>Institute of<br>Neurology | Author | Morphometric<br>analysis and data<br>acquisition |
| Yan Liu | UCL Queen square<br>Institute of<br>Neurology | Author | Digitisation of<br>images and<br>preparation for<br>analysis |
| Smriti Patodia | UCL Queen square<br>Institute of<br>Neurology | Author | Digitisation of<br>images and<br>preparation for<br>analysis |
| Zuzanna Michalak | UCL Queen square<br>Institute of<br>Neurology | Author | Preparation of<br>sections and tissue<br>acquisition from<br>brain banks |
| Sanjay Sisodiya | UCL Queen square<br>Institute of<br>Neurology | Author | Clinical data<br>acquisition, study<br>design and<br>interpretation and<br>review of the<br>manuscript |
| Maria Thom | UCL Queen square<br>Institute of<br>Neurology | Author | Study design, data<br>acquisition, data<br>analysis and<br>manuscript<br>preparation |

## List if Abbreviations

GCL: Granule cell layer
HMAL: Hippocampal malrotation
HS: Hippocampal sclerosis
PHG: parahippocampal gyrus
PM: post mortem
SUDEP: Sudden and unexpected death in epilepsy
TLE: temporal lobe epilepsy

## Notes

Disclosures: All authors report no disclosures.

Funding: UCL is part of the Center for SUDEP Research (CSR) and supported through the National Institute of Neurological Disorders And Stroke of the National Institutes of Health (Award Numbers neuropathology of SUDEP: 5U01NS090415 and SUDEP admin core grant: U01-NS090405). Epilepsy Society supports SMS, and through the Katy Baggott Foundation, supports the UCL Epilepsy Society Brain and Tissue Bank. This work was undertaken at UCLH/UCL who received a proportion of funding from the Department of Health’s NIHR Biomedical Research Centres funding scheme. ZM was funded by the European Union’s Seventh Framework Program (FP7/2007-2013) under grant agreement 602102 (EPITARGET).

